# Local topology, bifurcations and mutation hot-spots in proteins with SARS-CoV-2 spike protein as an example

**DOI:** 10.1101/2020.11.11.378828

**Authors:** Xubiao Peng, Antti J Niemi

## Abstract

Novel topological methods are introduced to protein research. The aim is to identify hot-spot sites where a bifurcation can change the local topology of the protein backbone. Since the shape of a protein is intimately related to its biological function, a mutation that takes place at such a bifurcation hot-spot has an enhanced capacity to change the protein’s biological function. The methodology applies to any protein but it is developed with the SARS-CoV-2 spike protein as a timely example. First, topological criteria are introduced to identify and classify potential mutation hot-spot sites along the protein backbone. Then, the expected outcome of a substitution mutation is estimated for a general class of hot-spots, by a comparative analysis of the backbone segments that surround the hot-spot sites. This analysis employs the statistics of commensurable amino acid fragments in the Protein Data Bank, in combination with general stereochemical considerations. It is observed that the notorious D614G substitution of the spike protein is a good example of such a mutation hot-spot. Several topologically similar examples are then analyzed in detail, some of them are even better candidates for a mutation hot-spot than D614G. The local topology of the recently observed N501Y mutation is also inspected, and it is found that this site is prone to a different kind of local topology changing bifurcation.

## Introduction

Topological methods are among the most versatile tools available to predict, model and analyze a wide range of Physics related phenomena, from theories of fundamental interactions to models of condensed matter and dynamical systems [1–4]. However, despite the apparently rich and intriguing topology of many biomolecules, thus far topological methods have been sparsely applied to biophysical problems. Among the notable exemptions are the analysis of enzyme action on DNA and the challenge to understand knots and their biological role in folded proteins [5].

Here a novel topological methodology, apparently with no previous physical applications, is introduced and developed for biophysical purposes. The methodology builds on inspection of bifurcations that can change the local topology of a protein backbone. Bifurcations are the common cause for qualitative changes in a physical system, bifurcation theory describes how a small change in an input parameter can cause a large scale change in the system [4, 6]. Accordingly it is proposed that in the case of a protein backbone, an amino acid site that is proximal to a bifurcation where the local topology can change, can be a mutation hot-spot where a single substitution can cause a large scale conformational change with potentially substantial biological effects: Even though such a mutation hot-spot is not necessarily more prone for a mutation than any other site along the backbone, when a mutation causes a bifurcation that changes the local topology it can also have a more profound biological impact. Thus the present methodology, that employs bifurcations to identify and classify potential mutation hot-spot sites, can have a value to a wide range of future investigations.

Indeed, the methodology that is developed here is very general. It is applicable to any protein structure, even though it is presented with the spike protein of the severe acute respiratory syndrome coronavirus 2 (SARS-CoV-2) as an example. The choice reflects the current urgency to understand the function of the virus that causes COVID-19, a global public health emergency that continues to spread across the world. Several studies of the SARS-CoV-2 virus have been published including investigations on the source of infection [7–11], the mechanism of transmission [12–16], and the structure and function of its various proteins [17–23]. These studies also detail the biophysical and biochemical properties of the spike protein, a transmembrane glycoprotein that assembles into a homo-trimer to cover the virion surface and gives the virus its distinctive crown-like look. For the present purposes the ensuing short introduction to the spike protein structure and function is sufficient:

The spike protein has three different stages. The prefusion stage, the intermediate stage, and the post-fusion stage. Before the fusion with a host cell takes place the spike protein is in the prefusion stage, and the present study focuses solely on the protein in this stage. The monomer structure of the prefusion stage is as follows: Starting from the N-terminal, first there is a short signal peptide. This is followed by two larger subunit, S1 and S2. The subunit S1 can recognize the host cell and bind to the receptor angiotensin-converting enzyme 2 (ACE2). The subunit S2 can bind to the membrane of the host cell, to mediate the fusion between the virus and the host cell. The Figure 1A identifies the major functional domains in these subunits in a monomeric spike protein [17, 18]: The S1 subunit consists of residues between the sites 14-685. It starts with a N-terminal domain (NTD) with residues 14-305 [25]. The NTD is followed by the receptor binding domain (RBD, residues 319-541) [26]. The junction segment between the S1 and S2 subunits includes several cleavage sites [27]. The S2 subunit that comprises the rest of the protein, contains the fusion peptide (FP, residues 788-806) followed by two heptapeptide repeat sequences (HR1, residues 912-984) and (HR2, residues 1163-1213), the transmembrane domain (TM, residues 1213-1237) and the cytoplasm domain tail (CT, residues 1237-1273) [24].

**Fig 1.**
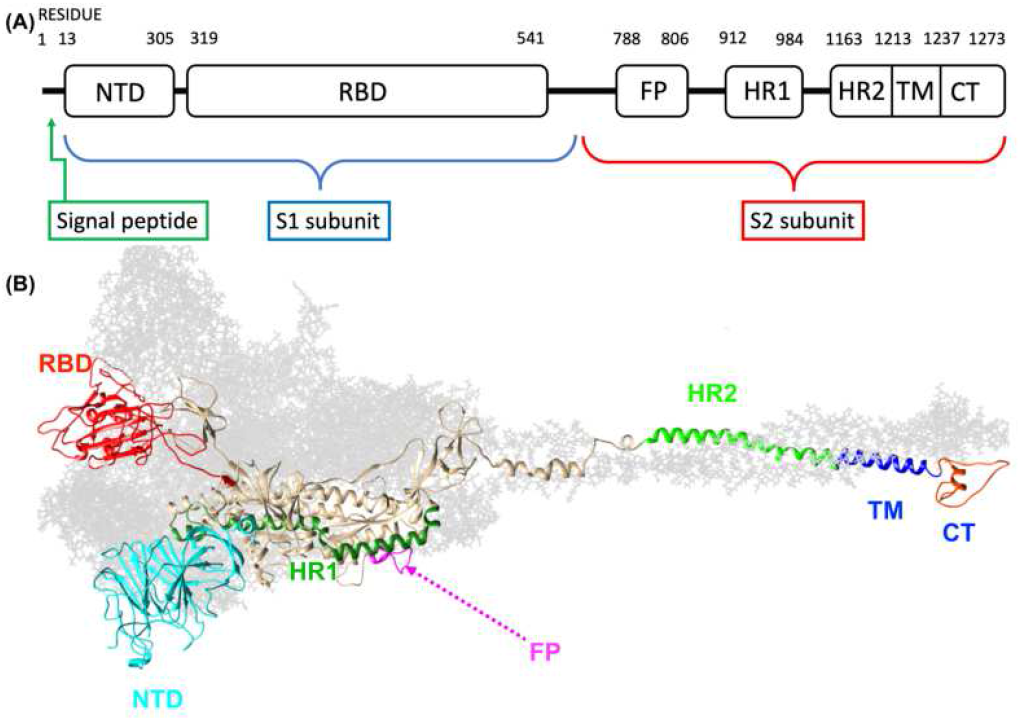
(A) Spike protein subunit S1 consists of the N-terminal domain NTD and the receptor binding domain RBD. The subunit S2 consists of the fusion peptide FP, two heptapeptide repeat sequences HR1 and HR2, the transmembrane domain TM and the cytoplasm domain tail CT. (B) A three-dimensional model of the full length closed stage spike protein, based on the PDB structure 6VXX (closed state) and adapted from [24].

In the prefusion stage the protein has two principal conformational states, called the closed state and the open state. The closed state is the native prefusion state of the spike protein, and a model of this closed stage is shown in Figure 1B. When the virus starts interacting with the host cell the spike protein transits from the closed state to the open state, with a major conformational change that takes place in the S1 subunit [19].

The spike protein has a pivotal role in processes that range from receptor recognition to viral attachment and entry into host cell. It is a major target in both vaccine research and therapeutic research that combat the SARS-CoV-2 virus. But the spike protein evolves and mutates continuously which makes it a demanding target for the development of antiviral inhibitors. Among the prominent examples of a mutation in the spike protein analyzed here, with documented epidemiological consequences, is the D→G substitution that occurred at its site 614, near the junction between the subunits S1 and S2. Apparently this mutation converted SARS-CoV-2 into a more transmissible form, and the D614G mutated virus now dominates in the global COVID-19 pandemic [30]. Another, more recent mutation that is also analyzed here, with similarly substantial epidemiological consequences that still remain to be fully understood, is the N→Y substitution that occurred at site 501 in the RDB domain [31]. Unfortunately, due to missing residues in the available data the even more recent substitutions that affect the sites 484 and 677 [32] can not yet be analyzed.

The goal of the present article is to develop general methodology that can help to understand the structural effects of these and other mutations, and to identify additional mutation hot-spot sites along the spike protein backbone, those where a single substitution mutation can have a large scale conformational effect. The methodology starts with a geometric scrutiny of a protein’s Cα backbone. For this, the Methods section first summarizes the known mathematical results on geometry and local topology of a smooth space curve. The focus is on the two curve specific bifurcations that can alter the local topology of a curve, these were introduced by Arnol’d [33–35] who called them the inflection point perestroika and the bi-flattening perestroika; see also [36, 37]. The Methods section then adapts these bifurcations to the case of a generic protein Cα backbone.

The Results section applies the general methodology to identify and classify mutation hot-spots. The SARS-CoV-2 spike protein is used as a timely example but any protein backbone can be analyzed in the same manner. For the spike protein, the Protein Data Bank structures 6VXX (closed state) 6VYB (open state) [19] and 6XS6 (D614G mutation) [38] are used. First, a general class of local topologies with a mutation hot-spot is identified. It consists of backbone segments that surround a site that is proximal to a flattening point: The D614G mutation occurred at such a hot-spot, thus the details of the methodology are worked out with the site 614 of the spike protein as an example. All similar hot-spot sites of the spike protein, with a site proximal to a flattening point, are tabulated in the currently available PDB structures. Additional examples are analyzed, including the identification of the likely amino acid substitution that may take place if a mutation occurs at such a hot-spot. Finally, the site of the N501Y substitution is identified as a different kind of mutation hot-spot, and analyzed as an additional example of the present approach.

## Methods

### Local topology of regular space curves

The bifurcations that can change the local topology of a space curve were introduced and analyzed by Arnol’d in a series of articles [33–35]; see also [36, 37]. This subsection summarizes the main results. The starting point is in the Frenet framing that governs the geometry of a regular, analytic space curve that is not a straight line [39]. To describe this framing, consider a parametric representation **x**(*s*) ∈ ℝ^3^ of a space curve, with *s* ∈ [0, *L*] the arc-length parameter and *L* its fixed length. The unit length tangent vector is

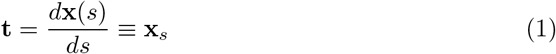

The unit length binormal vector is

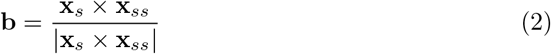

and the unit length normal vector is

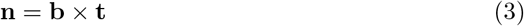

Together, the three vectors (**n**(*s*), **b**(*s*), **t**(*s*)) define the right-handed orthonormal Frenet frame at the regular point **x**(*s*) of the curve. They govern the geometry of the curve in terms of curvature *κ*(*s*)

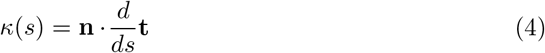

that describes how the curve bends on the osculating plane spanned by **t**(*s*) and **n**(*s*), and torsion *τ*(*s*)

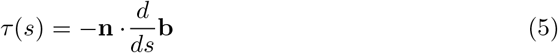

that measures how the curve deviates from this osculating plane.

The Frenet frames can be introduced whenever the curvature *κ*(*s*) is non-vanishing. In the specific limit where the curvature is very small in comparison to the torsion, but does not vanish

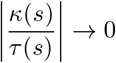

the frames obey

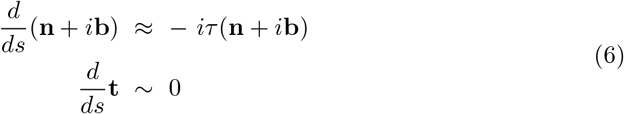

This limit, that not discussed in [33–37] but turns out to be relevant to protein backbones, describes a framed, straight line with framing that rotates around the line at a rate and direction that is determined by *τ*(*s*).

In the following it is assumed that the curve is open and its shape can change freely by local deformations, but the end points are always fixed. The shape changes are governed by the following rules [33–37]:

At a point where the curvature *κ*(*s*) vanishes, the Frenet frame can not be defined; here it is assumed that *κ*(*s*) has only isolated simple zeroes. The points where *κ*(*s*) = 0 are called the inflection points of the curve. A local deformation that retains the osculating plane can move an inflection point along the curve. But if a deformation lifts a point with *κ*(*s*) = 0 off the osculating plane the inflection point becomes removed. This implies that the co-dimension of an inflection point is two: An inflection point is not a local topological invariant of the curve, and a generic curve does not have any inflection points.

A point where the torsion vanishes *τ*(*s*) =0 is a flattening point. Unlike an inflection point, at least one single simple flattening point is generically present in a curve. A single simple flattening point is a local topological invariant that can not be removed by any continuous local deformation of the curve, it can only be moved along the curve.

The three vectors (**n**(*s*), **b**(*s*), **t**(*s*)) determine a framing of the curve, and either **b**(*s*) or **n**(*s*) or their linear combination can be chosen as the framing vector. The self-linking of the curve describes how it links with a nearby curve that is obtained by pushing points of the original curve along the framing vector. The self-linking number is a local topological invariant of the curve, in the absence of an inflection point the self-linking number can not change.

When the shape of a curve changes freely, an isolated inflection point generically occurs at some instance. When an inflection point appears the curve undergoes a bifurcation that is called an inflection point perestroika [33–35]. This bifurcation can change he number of flattening points: Since the torsion *τ*(*s*) changes its sign at a simple flattening point, and since the curvature *κ*(*s*) is generically not zero, an inflection point perestroika commonly changes the number of simple flattening points by two.

When the shape of a curve changes so that a pair of flattening points comes together they combine into a single bi-flattening point. A bi-flattening point can then be removed by a further, generic local deformation of the curve. Similarly, a bi-flattening point can first be created by a proper local deformation of a curve, and when the curve is further deformed the bi-flattening point can resolve into two separate simple flattening points. When either of these occur the curve undergoes a bifurcation that is called a bi-flattening perestroika [33–35].

Inflection point perestroika and bi-flattening perestroika are the only two bifurcations where the number of flattening points can change. Furthermore, according to [33–37]the number of flattening points and the self-linking number that isdetermined by the Frenet framing are the only two curve specific local topological invariants that can be assigned to a curve.

The number of flattening points and the self-linking number are independent topological invariants, but in the presence of an inflection point they can interfere with each other. For example, when a curve is deformed so that two simple flattening points become combined and disappear in a bi-flattening perestroika, the self-linking number in general does not change. However, if the bi-flattening perestroika occurs in conjunction of an inflection point perestroika the self-linking number can change: If the torsion is initially positive and two flattening points combine and disappear with the passage of an inflection point, the self-linking number increases by one. But if the torsion is initially negative the self-linking number decreases by one.

Note that the limiting case (6) is excluded in these analyses but it will become important in the sequel, in applications to protein backbones.

### The geometry and local topology of a protein C*α* backbone

The relations that are described in the previous subsection are valid for (thrice) continuously differentiable curves. In this subsection they are adapted to the case of a C*α* backbone that determines a piecewise linear polygonal chain, with C*α* atoms at the vertices.

Various shape changes are common in a biologically active protein. A polygonal chain such as the C*α* backbone can always be thought as a limiting case of a regular, analytic space curve. Thus the changes in the shape of the C*α* backbone are subject to same rules that govern the local topology of any regular space curve. In particular, the three essential shape deformations that can change the local topology *i.e.* discrete variants of inflection point perestroikas, bi-flattening perestroikas, and changes in a self-linking number should all have a profound role in physiological processes.

The discrete Frenet frame formalism is developed in [40]. The formalism describes the geometry of a piecewise linear chain with vertices **r**_*i*_ (*i* = 1,..., *N*) that in the case of a protein backbone are the space coordinates of the C*α* atoms. A line segment defines the unit tangent vector

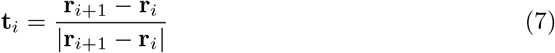

It points from the center of the *i^th^* C*α* atom towards the center of the (*i* + 1)^st^ C*α* atom. The unit binormal vector is

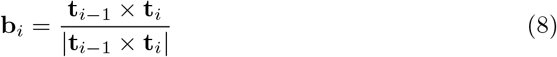

and the unit normal vector is

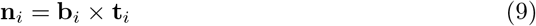

In the case of a protein these two vectors are virtual in that they do not point towards any particular atom. The orthonormal triplet (**n**_*i*_, **b**_*i*_, **t**_*i*_) defines a discrete version of the Frenet frames (1)-(3) at each position **r**_*i*_ along the chain. In lieu of the curvature *κ*(*s*) and the torsion *τ*(*s*) there are now their discrete versions, the bond angles *κ*_*i*_ and the torsion angles *τ*_*i*_. The values of these angles can be computed from the discrete Frenet frames. The bond angles are

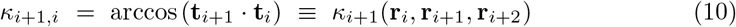

and the torsion angles are

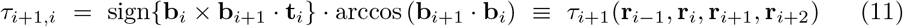

It is notable that the value of the bond angle *κ*_*i*_ is evaluated from three, and the value of the torsion angle *τ*_*i*_ is evaluated from four consecutive vertices.

Conversely, when the values of the bond and torsion angles are all known, the discrete Frenet equation [40]

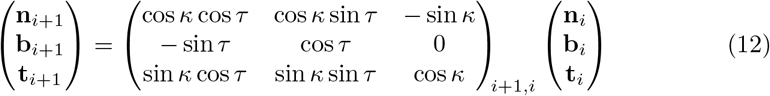

computes the Frenet frame at the vertex **r**_*i*+*i*_ from the frame at the preceding vertex **r**_*i*_. In a continuum limit [40] the discrete Frenet equation becomes the continuum Frenet equation [39].

The fundamental range of the bond angles is κ_*i*_ ∈ [0, π] and in the case of the torsion angles *τ*_*i*_ ∈ [–*π,π*). For visualization purposes, the bond angles κ_*i*_ can be identified with the latitude angle of a two-sphere that is centered at the *i^th^* C*α* atom; the north pole coincides with the inflection point κ_*i*_ = 0. The torsion angles *τ_i_* ∈ [–*π, π*) correspond to the longitudinal angle on the sphere, the value increases in the counterclockwise direction around the tangent vector and the value *τ*_*i*_ = 0 of flattening points coincides with the great semi circle that starts from north pole and passes through the tip of the normal vector **n** to the south pole. The sphere can be stereographically projected onto the complex (*x, y*) plane. A projection from the south pole is

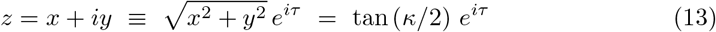

as shown in figure 2: The north pole *i.e.* the point of inflection with *κ* = 0 becomes mapped to the origin (*x, y*)=(0, 0) and the south pole *κ* = *π* is sent to infinity.

**Fig 2.**
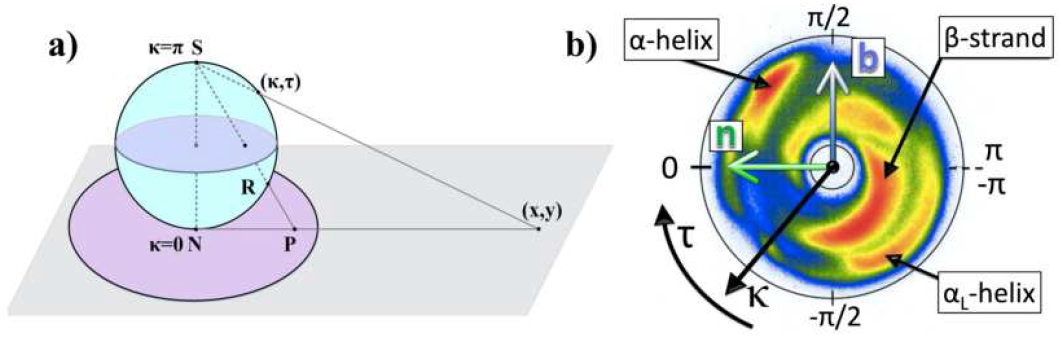
a) Stereographic projection of two sphere from the north pole (N) with latitude *κ* and longitude *τ*. b) The stereographically projected Frenet frame map of backbone C*α* atoms, with major secondary structures identified. Also shown is the directions of the Frenet frame normal vector **n** and bi-normal vector **b**; the vector **t** points upwards from the figure. Colour coding corresponds to the number of PDB entries with red large, blue small and white none.

The C*α* backbone can be visualized on the stereographically projected two sphere as follows [41]: At each C*α* atom, introduce the corresponding discrete Frenet frames (7)-(9). The base of the *i^th^* tangent vector **t**_*i*_ that is located at the position **r**_*i*_ of the *i^th^* C*α* atom coincides with the centre of a two-sphere with the vector **t**_*i*_ pointing towards its north pole. Now, translate the sphere from the location of the *i^th^* C*α* atom to the location of the (*i* + 1)^*th*^ C*α* atom, without any rotation of the sphere with respect to the *i^th^* Frenet frames. Identify the direction of **t**_*i*+1_, i.e. the direction towards the C*α* atom at site **r**_*i*+2_ from the site **r**_*i*+1_, on the surface of the sphere in terms of the ensuing spherical coordinates (*κ_i_, τ_i_*). When this construction is repeated for all the protein structures in Protein Data Bank that have been measured with better than 2.0 Å resolution, the result can be summarized by the map that is shown in figure 2 b). The color intensity correlates directly with the statistical distribution of the (*κ_i_,τ_i_*); red is large, blue is small and white is none. The map describes the direction of the C*α* carbon at **r**_*i*+2_ as it is seen at the vertex **r**_*i*+1_, in terms of the Frenet frames at **r**_*i*_.

Approximatively, the statistical distribution in figure 2 b) is concentrated within an annulus that corresponds to the latitude angle values (in radians)*κ* = 0.57 and *κ* = 1.82 shown in the Figure. The exterior of the annulus is a sterically excluded region. The entire interior is sterically allowed, but there are very few entries in this region. The four major secondary structure regions, *α*-helices, *β*-strands, left-handed *α*-helices and loops, are identified according to their PDB classification. For example, (*κ, τ*) values (in radians) for which

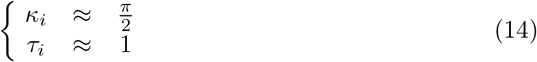

describes a right-handed *α*-helix, and values for which

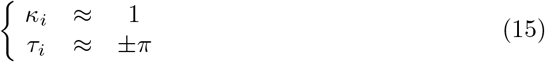

describes a *β*-strand.

In the case of a regular space curve both *κ*(*s*) and τ(*s*) are smooth functions and the inflection points and the flattening points are easily identified as the points where *κ*(*s*) = 0 and *τ*(*s*) =0. In a crystallographic protein structure where the C*α* positions are experimentally determined, an inflection point is detectable as a very small value of the bond angle *κ*_*i*_ at the proximal C*α* vertex. Similarly, the presence of a simple flattening point can be deduced from a very small torsion angle value at the proximal vertex, accompanied by a sign change in *τ* between two neighboring vertices; if the sign of *τ* does not change the proximal vertex has the character of a bi-flattening point. Accordingly, when searching for C*α* atoms where essential shape changes such as inflection point or bi-flattening perestroikas can take place, the natural points to start are the neighborhoods of vertices where either *κ*_*i*_ ≈ 0 or *τ*_*i*_ ≈ 0. These are the likely locations where a small change in the shape of the backbone can change its local topology, with a potentially substantial change in the protein’s biological function.

From Figure 2 b) one observes that inflection points *i.e.* small *κ*_*i*_ values are extremely rare in crystallographic protein structures. Indeed, a generic space curve does not have any inflection points. At the same time general arguments state that generically at least one flattening point can be expected to be present. As shown in Figure 2 b) flattening points where *τ*_*i*_ ≈ 0 do appear even though they are relatively rare in protein structures. Moreover, it is observed from the Figure that at a flattening point the bond angle values are mostly either *κ*_*i*_ ≈ 1 or *κ*_*i*_ ≈ *π*/2.

Since the torsion angles are defined *mod*(2*π*) as implied in (15), in the case of discrete Frenet frames there is an additional structure: There is the line *τ*_*i*_ = ±*π* in Figure 2 b) where the torsion angle has a 2*π* discontinuity, hence it can change sign by crossing the line. This multivaluedness is absent in regular space curves, with *τ*(*s*) a single-valued continuous function. But the limits *τ_i_* → ±*π* can be thought of as the small curvature and large torsion limits of the equations (6). Notably, *κ_i_* = 1 and *τ_i_* → ±*π* corresponds to an ideal, straight *β*-strand. Therefore a point on the line *τ_i_* → ±*π* (or in its immediate vicinity) will be called a *β*-flattening point in the sequel.

The multi-valuedness of the torsion angle *τ*_*i*_ affects the local topological invariants, in the case of a discrete chain. This is exemplified in the Figures 3. These Figures depict the three characteristic examples of a protein loop topology, when the loop interpolates between two right-handed *α*-helices; similar considerations apply when the interpolation is between any two generic points.

**Fig 3.**
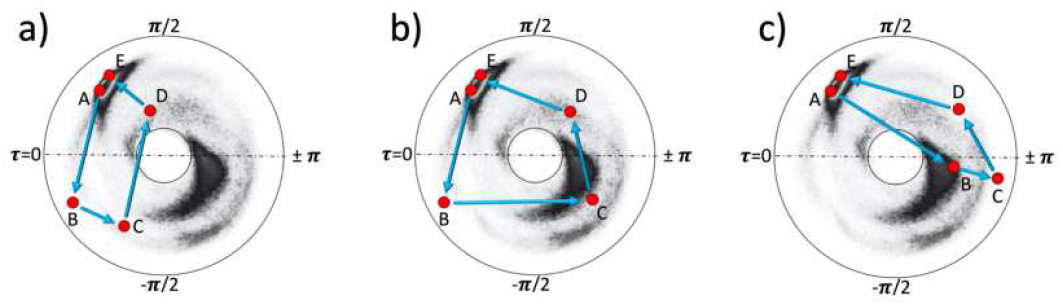
Different local topologies for a chain with five vertices. a) With two crossings of the flattening point line *τ* = 0. b) With one crossing of the flattening point line *τ* = 0 and one crossing of the line of *β*-flattening points *τ* = ±*π*. c) With two crossings of the line of *β*-flattening points *τ* = ±*π*. In both a) and c) the folding index Eq(16) vanishes but in b) the folding index has value −1 since the trajectory encircles the disk center once in counterclockwise direction.

For clarity: Whenever a Figure similar to those in Figures 3 is drawn in the sequel, the two neighboring vertices i.e. values (*κ*_*i*_,*τ*_*i*_) and (*κ*_*i*+1_,*τ*_*i*+1_) are always to be connected by a virtual segment that is a straight line on the disk. Even though in reality those values of (*κ, τ*) that are on the straight line may not correspond to any actual atomic position along the piecewise linear C*α* backbone. At the level of local topology, this has no effect.

In all Figures 3, for convenience the initial points (A) and the final points (E) are fixed and located near the *α*-helical region where the torsion angle has a positive value close to *τ* ≈ 1 but this can be changed with no effect. At the points (B) and (C) the torsion angles are always negative in the Figures 3. At the point (D) the torsion angle is positive.

In the Figure 3 a) the chain proceeds from point (A) to point (B) by crossing the line of flattening points *τ* = 0. It then continues to point (C). From there it proceeds and crosses the line of flattening points a second time, to arrive at point (D) where the torsion angle returns to a positive value. The chain then continues and ends at point (E). In this case the local topology is fully analogous to that of a regular curve. In particular, the two flattening points along the chain can be removed by a bi-flattening perestroika that lifts both points (B) and (C) above the line *τ* = 0 to positive *τ_i_* values, without the chain crossing the inflection point *κ* = 0 at the center of the disk.

In the Figure 3 b) the chain proceeds to the point (B), again by crossing the line of flattening points *τ* = 0. From there it proceeds to point (C) that is located near the *β*-stranded region. The chain then crosses the line of *β*-flattening points *τ* = ±*π* as it proceeds to point (D) where the torsion angle is positive. The chain finally ends at point (E) in the *α*-helical region. In this case there is only one flattening point along the chain, since the second time the torsion angle changes its sign at a *β*-flattening point. The chain encircles the inflection point once in the counterclockwise direction. Neither the flattening point nor the *β*-flattening point can be removed without encountering an inflection point perestroika along the chain.

The Figure 3 b) motivates to introduce a winding number termed folding index *Ind_f_* [42] for a backbone chain segment between sites *n*_1_ and *n*_2_. The folding index classifies loop structures and entire folded proteins by counting the number of times the chain encircles the inflection point. Its value can be obtained from the equation

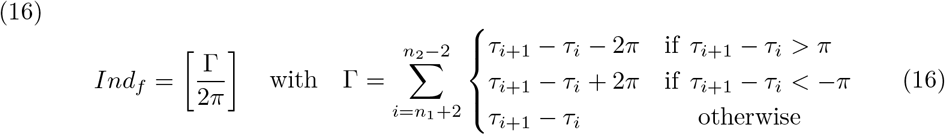

with [*x*] the integer part of *x*. Here Γ is the total rotation angle (in radians) that the chain makes around the inflection point in Figures such as 3. The folding index is positive when the rotation is clockwise, and negative when the rotation is counterclockwise. In the Figure 3 b) the folding index has the value *Ind_f_* = –1 since the chain encircles the inflection point once in counterclockwise direction.

In the Figure 3 c) the torsion angle changes its sign between points (A) and (B) and between points (C) and (D). Now the sign change occurs at *β*-flattening points. The general rules of local topology for the sign change at a *β*-flattening point are like those at a flattening point. In particular, a pair of *β*-flattening points can either be created or removed in a bifurcation called *β*-flattening perestroika that is akin a bi-flattening perestroika. In the Figure 3 c) a *β*-flattening perestroika occurs if the vertices (B) and (C) are both moved upwards across *τ* = ±*π* line without the chain crossing the inflection point *κ* = 0 at the center of the disk.

The folding index is a local topological invariant. For example, in Figure 3 b) its value does not change unless the chain is deformed so that either the flattening point between (A) and (B) or the *β*-flattening point between (C) and (D) becomes removed by an inflection point perestroika, that either converts the *β*-flattening point into a flattening point by a deformation that takes the chain in Figure 3 b) into the chain in Figure 3 a) or converts the flattening point into a *β*-flattening point by a deformation that takes the chain in Figure 3 b) into the chain in Figure 3 c). In both cases the final folding index vanishes.

The three examples in Figures 3 summarize all essential aspects of local topology that are encountered in the case of a discrete chain. In particular, the line of flattening points and the line of *β*-flattening points have a very similar character in terms of local topology. They can be interchanged by an inflection point perestroika that also changes the folding index.

Finally, it is noted that the Figure 2 b) is akin the Newman projection of stereochemistry. The vector **t**_*i*_ which is denoted by the red dot at the center of the figure, points along the backbone from the proximal C*α* at **r**_*i*_ towards the distal C*α* at **r**_*i*+1_, and the colour intensity displays the statistical distribution of the **r**_*i*+2_ direction. Moreover, unlike the Ramachandran map the figure 2 b) provides non-local information on the backbone geometry. The Ramachandran map can only provide localized information in the immediate vicinity of a single C*α* carbon, but the information content in the figure 2 b) map extends over several peptide units. As shown in [43], the C*α* backbone bond and torsion angles (*κ*_*i*_,*τ*_*i*_) are sufficient to reconstruct the entire backbone, but the Ramachandran angles are not.

## Results

It turns out that the spike protein site 614 of the notorious D→G substitution is a good example of a site that is proximal to a flattening point. This motivates to select the flattening points of the spike protein as a starting point for presenting the general methodology; the methodology is quite independent of this choice. A detailed analysis of other potential bifurcation structures, those that engage a site that is proximal to an inflection point or a *β*-flattening point, is presented elsewhere. However, in the sequel it is also proposed that the recently observed N501Y mutation can provide an example that engages the inflection point.

First, the sites along the SARS-CoV-2 spike protein C*α* backbone that are proximal to a flattening point are classified. Their local neighborhoods are then inspected by comparisons with Figures 3, to investigate the potential bifurcations.

The Table 1 lists all those SARS-CoV-2 spike protein C*α*-sites that are proximal to a flattening point in the PDB structures 6VXX (closed state) and 6VYB (open state). Here a torsion angle value is determined to be proximal to a flattening point when |*τ*_*i*_| < 0.2. In terms of distance, this is less than the radius of a carbon atom. The histogram in Figure 4 shows the distribution of the flattening point sites in relation to spike protein backbone. There are relatively many entries in the NTD and RBD domains, and in the fusion core between the HR1 and HR2 domains. Notably the number of proximal sites is also different in the closed and open states of the spike protein, there are more proximal sites in the open state. Thus a transition between the two states involves changes in local topology that affect in particular the NTD and RBD domains, and the HR1-HR2 junction.

**Table 1.**
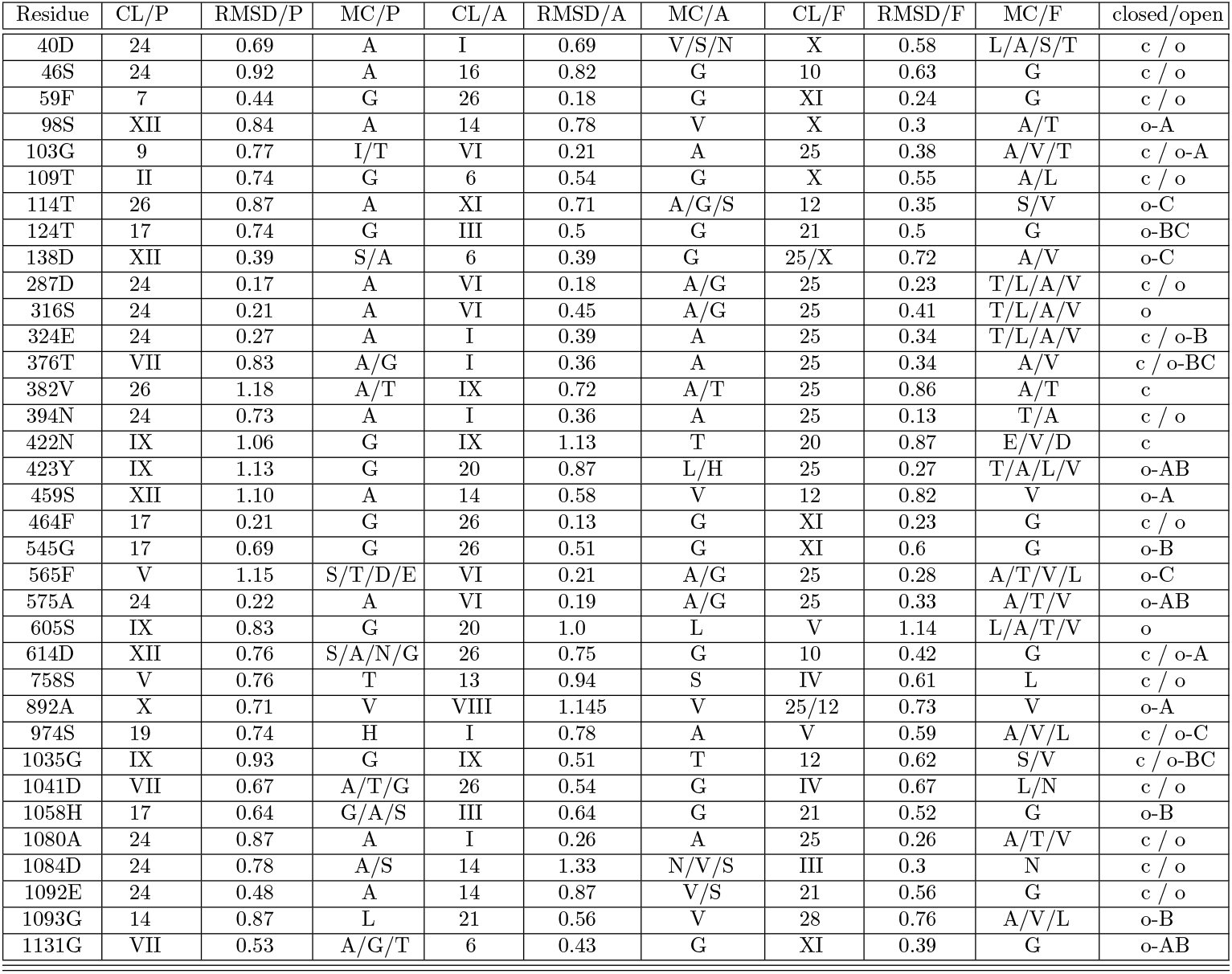
Flattening points in spike protein PDB structures 6VXX and 6VYB. Columns are organized as follows: Residue number with present amino acid. CL stands for cluster with smallest RMSD (in Å); clusters are taken from [44]. MC stands for most common mutagenic amino acid substitutions in the cluster. P stands for preceding, A stands for adjacent and F stands for following fragment. Closed (c) and open (o) denotes the state where flattening point is observed. A,B,C denotes the monomer of spike protein.

**Fig 4.**
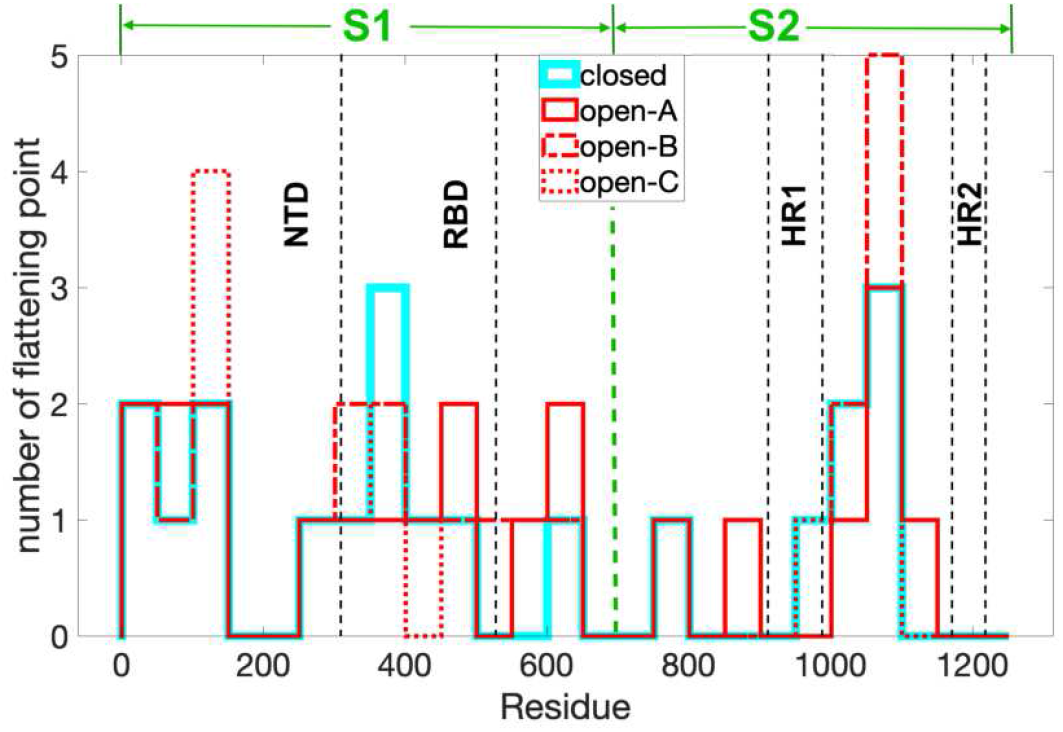
The histogram shows the distribution of sites that are proximal to a flattening point along the spike protein backbone, in blocks of 40 sites. The proximal sites accumulate in the NTD and RBD domains, and in the fusion core between the HR1 and HR2 domains. The PDB structures are 6VXX for closed state and 6VYB for open state.

In the case of a bond angle, here the value is considered small when *κ_*i*_* < 0.5. General arguments state that inflection points are not generic, and Figure 2 b) shows that there are indeed very few sites that are close to an inflection point. For the spike protein the smallest value is *κ_i_* = 0.39 and it is located at the site 103 that also appears in Table 1.

The local geometry of all the individual residues that are proximal to a flattening point has been investigated and the potential for a mutation hot-spot at a residue that is proximal to a flattening point has been estimated, by comparison to Figures 3. For this, a combination of statistical analysis and stereochemical constraints has been utilized. The Table 1 also summarizes these findings.

The statistical analysis employs the classification scheme of Protein Data Bank structures described in [44]. This scheme decomposes a C*α* backbone into fragments that consist of backbone segments with six successive sites; in accordance with Eq. (10), (11) a fragment with six sites determines three pairs of bond and torsion angles. In [44] the fragments that appear in high resolution PDB structures have been organized into disjoint clusters. To assign a cluster to a fragment, there must be at least one other fragment in the same cluster within a prescribed RMS cut-off distance;in [44] the cut-off is 0.2Å. Two clusters are then disjoint, when the RMSD between any fragment in the first cluster and any fragment in the second cluster exceeds this RMS cut-off distance. It was found that around 38% of protein loops in the high resolution PDB structures can be decomposed into fragments that belong to twelve disjoint clusters, labeled I-XII in [44]. When fragments from an additional set of 30 disjoint clusters are included, the coverage increases to ~ 52% [44].

In the Table 1 both cluster sets I-XII and 1-30 appear; the notation of [44] is used throughout in the following. Beyond these two sets, the clusters become increasingly smaller, and in the present study those smaller clusters have not been considered. The somewhat low resolution of the available spike protein structures in comparison to the very high resolution structures used in [44] does not justify a more detailed scrutiny.

The clusters in the Table 1 have been identified as follows: A pair of bond and torsion angles (*κ_i_, τ_i_*) at the *i^th^* site of the spike protein that is a proximal site to a flattening point can be assigned to three different clusters. The first cluster describes the bond and torsion angles for the sites (*i* – 2, *i* – 1, *i*); this cluster is labelled P (for Preceding) in the Table 1. The second cluster that is labelled A (for Adjacent) in the Table 1 describes the angles for sites (*i* – 1, *i, i* + 1). The third cluster is labelled F (for Following) and it describes the angles at spike protein sites (*i, i* + 1, *i* + 2). The cluster that provides the best match to the spike protein is listed in the Table 1 and it is determined as follows: Let (*x_a_, y_a_, z_a_*) denote the six space coordinates of a segment that corresponds to three consecutive pairs of (*κ, τ*) values in the spike protein. Let (*x_k,a_, y_k,a_, z_k,a_*) be the corresponding six space coordinates of a *k^th^* fragment in each of the clusters of [44]. The best matching cluster in Table 1 is the one that contains a fragment with the minimal root-mean-square distance (RMSD) to the given spike protein segment (in units of Ångström):

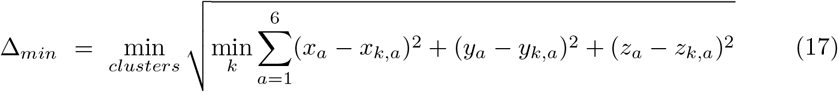

Once the best matching cluster is determined, the corresponding statistical distribution of amino acids found in [44] is used to identify those amino acids that are most probable to appear at the *i^th^* site of the spike protein, in case a mutation occurs. If the size of this statistically most probable amino acid is smaller than the size of the present amino acid at the *i^th^* site, a substitution by mutation is considered to be sterically possible. But if the size is larger, a substitution likely requires an extended rearrangement of the spike protein conformation and this can be energetically costly. The Table 1 lists the most probable substitutions that are predicted in this manner.

### Example: The D614G mutation of spike protein

The Table 1 identifies the site 614 of the spike protein, where the D→G substitution has occurred, as a site that is proximal to a flattening point. Thus, this site serves as a good example to describe the present methodology. The local topology and its potential bifurcations can be analyzed using figures such as Figure 5. Each of the three Figures depicts the three bond and torsion angle pairs for the three backbone segments (P, A and F respectively) that include the angles of the site 614.

**Fig 5.**
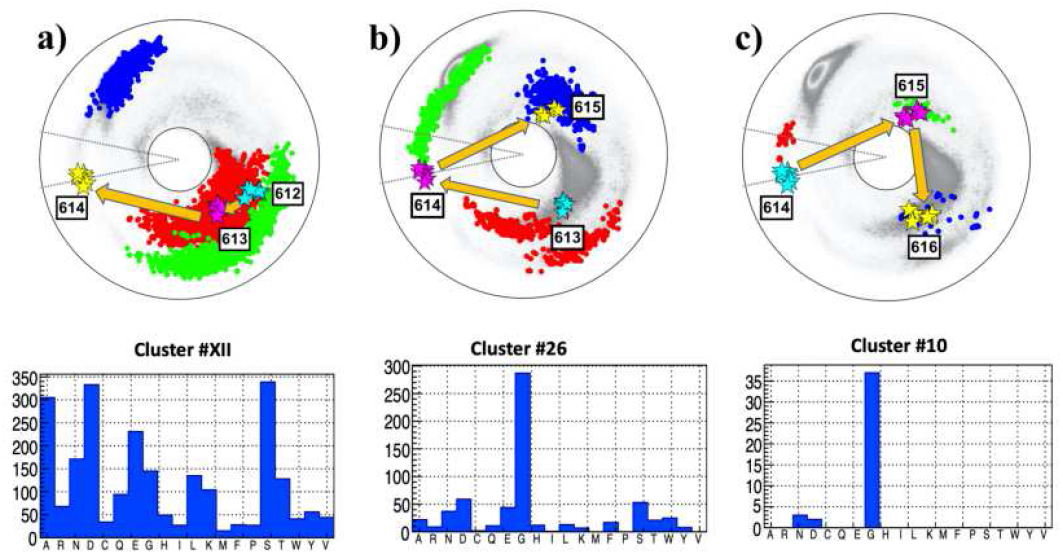
The (*κ_i_,τ_i_*) spectrum for *i* = 612 – 616; this engages sites 610 – 617 along the C*α* backbone. a) The spectrum for preceding sites P:(612, 613, 614), in the background of the best matching cluster #XII in [44]. The most common amino acids in the cluster are S, D, A, E, N, G. b) The spectrum for adjacent sites A:(613, 614, 615), in the background of the best matching cluster #26 in [44]. The most common amino acid in the cluster is G. c) The spectrum for following sites F:(614, 615, 616), in the background of the best matching cluster #10 in [44]. The most common amino acid in the cluster is G.

The legend is as follows: In figures such as Figure 5, the vertices that mark the C*α* sites of spike protein are always identified by stars. These stars are color-coded and organized according to increasing site number following the Cyan-Magenta-Yellow (CMY) color table. The dotted background comprises the C*α* sites of all the fragments in the corresponding best matching cluster. The background sites are always ordered similarly using the Red-Green-Blue (RGB) color table. The adjacent histogram displays the amino acid distribution in the best matching cluster, using the data that is obtained from [44].

The Figure 5 a) shows the bond and torsion angles for the sites P:(612,613,614) of the spike protein. The best matching preceding (P) cluster is also shown, it is the cluster #XII of [44]. From Table 1 the RMSD (17) between the spike protein segment and the cluster has the value Δ_*min*_ ≈ 0.76Å. This is clearly larger than the 0.2 Å cut-off used in [44]. Nevertheless, the cluster is included in the analysis since the resolution at which the spike protein PDB structures (6VXX, 6VYB and 6XS6) have been measured is also clearly larger than the resolution of the structures used in [44]. This justifies that despite the relatively large value of Δ_*min*_ the present analysis proceeds with cluster #XII, keeping in mind that the Δ_*min*_ is not very small.

The statistical analysis [44] shown in Figure 5 a) proposes that the most probable mutations at site 614D are to S, A, N and G; the amino acid E can be excluded on sterical grounds since it has a larger size than D.

The best matching cluster for the adjacent sites A:(613,614,615) shown in Figure 5 b) is the cluster #26 of [44], with Δ_*min*_ ≈ 0.75Å which is again relatively large. The statistical analysis [44] now proposes that the most probable mutation at site 614 is a D→G substitution; the probability for any other amino acid substitution is very low. In particular the probability for D itself is low, suggesting instability.

Finally, the best matching cluster for the following sites F:(614,615,616) shown in Figure 5 c) is #10. The RMSD has now a somewhat lower value Δ_*min*_ ≈ 0.41Å. The statistical analysis [44] proposes that the most probable mutation at site 614 is again D→G substitution. The probability for any other substitution is very low. In particular the probability for D itself is low, again suggesting instability of the residue.

The combined spike protein chain shown in the three Figures 5, starting from site 612 in 5 a) and ending at 616 in 5 c), encircles the inflection point once in clockwise direction. Thus the folding index has value +1 and, except for the direction and the location of the end points, the topology of the trajectory is similar to that in Figure 3 b). In particular, both a flattening point and a *β*-flattening point occur at neighboring segments along the chain. This proposes that the site 614 is a potential mutation hot-spot, prone to a change in the local topology by an inflection point bifurcation. The mutation can cause a bifurcation that can change the local topology from that resembling Figure 3 b) to one that resembles either Figure 3 a) with a bi-flattening point or Figure 3 c) with a pair of *β*-flattening points. Since G is the only amino acid that consistently appears in all three clusters and since there is no obvious steric hindrance for a D→G mutation, the prediction of the present analysis is that a D→G mutation is probable at the site 614 of spike protein, if a mutation indeed occurs; this is the notorious D614G mutation that has already been observed.

A comparison of the three PDB structures 6VXX, 6VYB with 6XS6 shows that apparently the mutation has not caused any change in local topology, at least according to available structures using the available resolution. But since the site 614 is located near the junction between S1 and S2 subunits, the D→G mutation in combination with the proximity of a flattening point has probably increased the chain flexibility in the junction segment so that it is now more prone to a cleavage with enhanced infectiousness as a consequence.

### A survey of flattening point hot-spots in spike protein

The Table 1 proposes that there is quite a large number of potential mutation hot-spots in the spike protein that can be similar to 614D, with a proximal flattening point. In the sequel a selection of these sites is analyzed. The examples are representative, but not necessarily the most probable hot-spot sites. There are three examples from the NTD domain, with site numbers 59, 103 and 287. The example at the site 103 is added since this is the site with the lowest bond angle value along the entire spike protein backbone. There is one example that is located in the junction between NTD and RBD domains, with site number 316. There is one example in the RBD domain, with site number 464. An example from the fusion core between HR1 and HR2 domains with site number 1080 is also presented. Finally, the local topology around the site 501 where the N→Y substitution has recently been observed [31] is also analyzed and its bifurcation potential is investigated.

### The residue 59F

The Figures 6 show the neighborhood of the site 59F, located in the NTD domain of subunit S1. The Figures reveal that the topology of the trajectory from site 57 to 61 is very similar to that in the case of site 614, shown in Figures 5: There is a residue that is proximal to a flattening point and right after it there is a residue that is proximal to a *β*-flattening point. The folding index has value +1 since the trajectory encircles the inflection point in clockwise direction. Thus, as in the case of 614, the site 59 is prone to an inflection point bifurcation such as those described in Figures 3. The RMSD values (17) are all quite small, indeed clearly smaller than in the case of 614D, so that the three clusters that are identified in the Figures 6 are a very good match. The statistical analysis of all three clusters show that G has a very high probability at the site 59; both S and D have some propensity albeit much smaller than G while the probability of the existing amino acid F is very small. Thus the site 59 is a very good candidate for a F→G mutation hot-spot.

**Fig 6.**
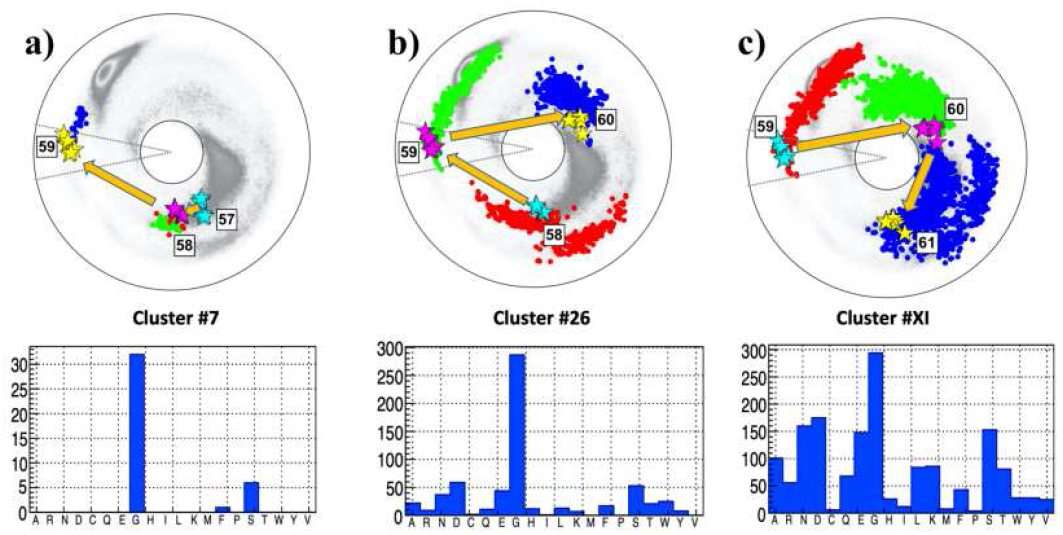
The (*κ_i_, τ_i_*) spectrum for *i* = 57 – 61; this engages sites 55 – 62 along the C*α* backbone. a) The spectrum for P:(57, 58, 59), in the background of the best matching cluster #7 in [44]. b) The spectrum for A:(58, 59, 60), in the background of the best matching cluster #26 in [44]. c) The spectrum for F:(59, 60, 61), in the background of the best matching cluster #XI in [44]. The most common amino acid in all three clusters is G.

### The case of 103

The Figures 7 show the neighborhood of the site 103G, located in the NTD domain of subunit S1. Here the situation is somewhat exceptional, since the residue 103 is already the smallest amino acid G.

**Fig 7.**
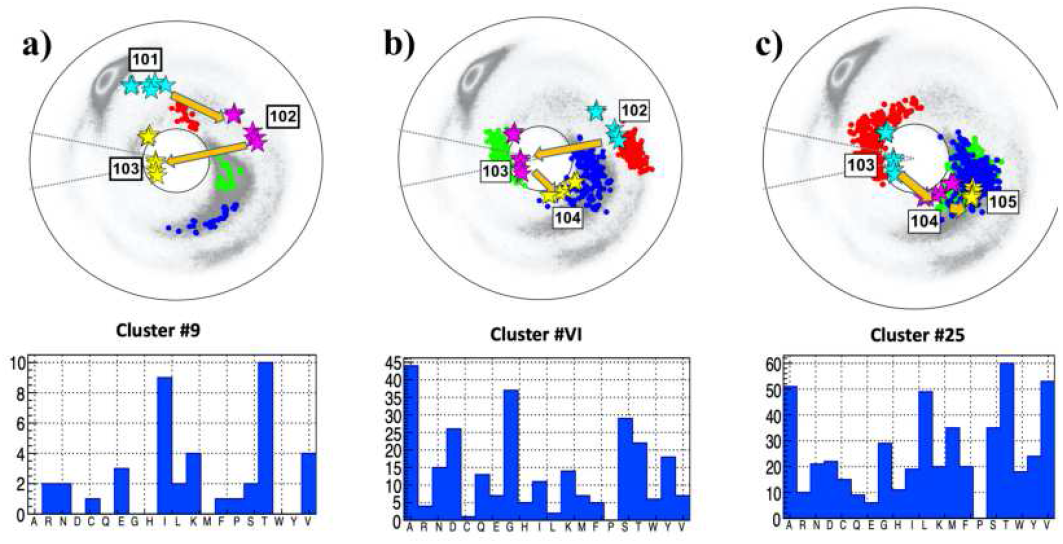
The (*κ_i_,τ_i_*) spectrum for *i* = 101 – 105; this engages sites 99 – 106 along the C*α* backbone. Panel a) The spectrum for P:(101,102,103), in the background of the best matching cluster #7 in [44]. The most common amino acids in the cluster is T, followed by I. Panel b) The spectrum for A:(102,103,104), in the background of the best matching cluster #26 in [44]. The most common amino acid in the cluster is A, followed by G. Panel c) The spectrum for F:(103,104,105), in the background of the best matching cluster #XI in [44]. The most common amino acid in the cluster are T and L, followed by A and V.

The chain between sites 101 and 105 is shown in Figures 7. It has one vertex near a *β*-flattening point at 102. This is immediately followed by the vertex 103 that is proximal both to a flattening point and to the inflection point. The following vertex 104 has also a very small bond angle value. The overall shape of the trajectory suggests a mutation hot-spot with inflection point perestroika that converts the site 103 from a vertex that is proximal to the flattening point into a vertex that is proximal to a *β*-flattening point. It is also plausible that there has been a recent mutation with ensuing inflection point perestroika, that has converted the local topology by moving the vertex 103 from the vicinity of a *β*-flattening point to the vicinity of a flattening point.

The statistical analysis shows that A, which is the smallest amino acids after G, has the highest propensity in the case of the adjacent cluster, shown in Figure 7 b). The propensity of A is also larger than that of G in the following cluster shown in Figure 7 c). Both clusters have also small RMSD value. On the other hand, in the histogram of the preceding cluster shown in Figure 7 a) the amino acid A is absent.

However, RMSD value is not very small, and the Figure also shows that cluster #9 can not be good match to the spike protein segment P:(101,102,103): The distance between the observed *τ* value at the site 103 deviates from the *τ*-values in the cluster by some 150 degrees. Thus the conclusion is that the cluster #9 should be used with care, for a mutation outcome prediction.

The Figure 8 a) shows the present 3D spike protein structure in the neighborhood of the site 103. In the Figure 8 b) the amino acid G has been replaced by A; the effect of this substitution is estimated using a crude energetic analysis with Chimera [45]. An inspection of the interatomic distances show that A can be substituted for G without encountering steric clashes. But in the case of the other amino acids T, V and L that also have a high propensity in the cluster of Figure 7 c), a substitution using Chimera leads to steric clashes.

**Fig 8.**
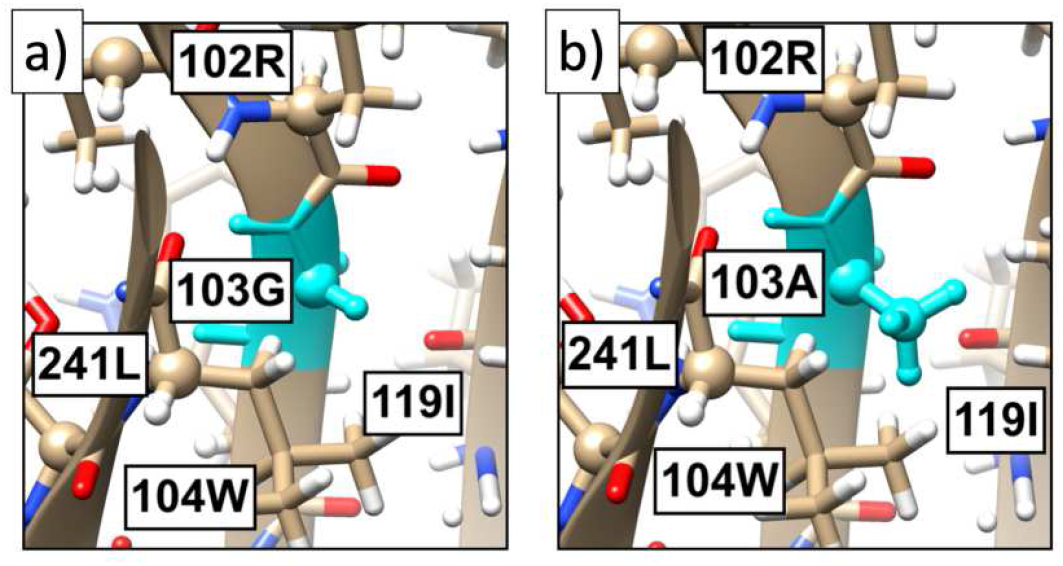
a) The present all-atom structure of the spike protein in the neighborhood of site 103. b) The substitution of A in place of 103G, as predicted by Chimera [45]. There are no apparent steric hinders for the substitution.

The conclusion of the present analysis is that the site 103G is a potential G→A mutation hot-spot.

### The case of 287D

The Figures 9 show the neighborhood of the site 287D, located in the NTD domain of subunit S1. The site is both preceded and followed by a site that is proximal to a *β*-flattening point. The folding index has value −1 since the chain encircles the inflection point in counterclockwise direction. But since 286 and 288 are both very close to a *β*-flattening point the chain passes back-and-forth very close to the inflection point.: The links 286-287 and 287-288 have a very similar topology to the links 102-103 and 103-104 in Figures 7.

**Fig 9.**
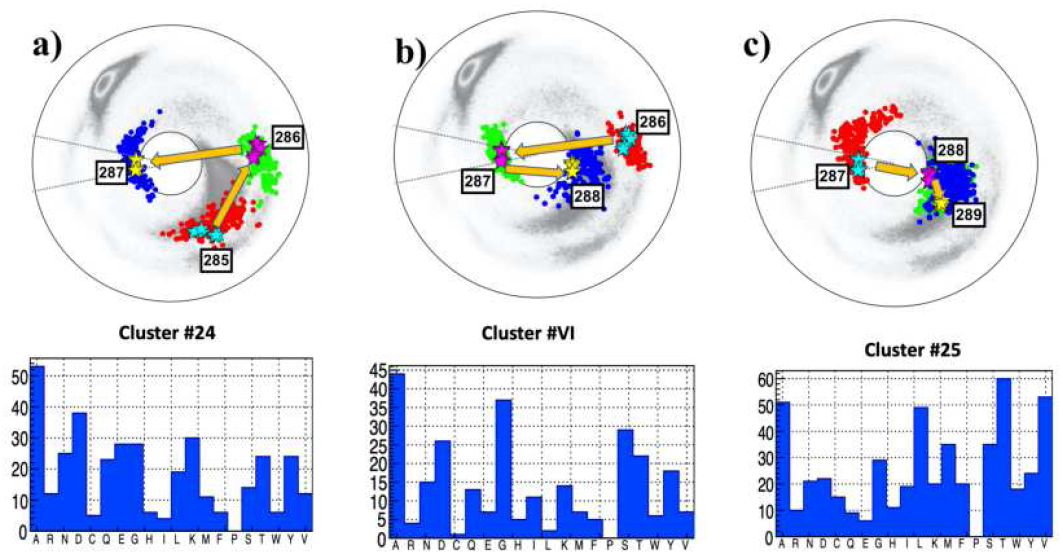
The (*κ_i_,τ_i_*) spectrum for *i* = 285 – 289; this engages sites 283 – 290 along the C*α* backbone. Panel a) The spectrum for P:(285, 286, 287), in the background of the best matching cluster #24 in [44]. The most common amino acids in the cluster is A, followed by D. Panel b) The spectrum for A:(286, 287, 288), in the background of the best matching cluster #VI in [44]. The most common amino acid in the cluster is A, followed by G. Panel c) The spectrum for F:(287, 288, 289), in the background of the best matching cluster #25 in [44]. The most common amino acid in the cluster are T and V, followed by A and L.

The RMSD values (17) are all very small so that the three clusters identified in the Figures 9 are an excellent match. The statistical analysis in all three clusters show that A has a very high probability at the site 287; both T and V are also likely substitutions in the following cluster, and G has also propensity in the adjacent cluster. But D that is now located at the site, is not very prominent in any of the clusters. Thus the site 287 is a very good candidate for a D→A mutation hot-spot.

### The case of 316S

The site 316S is located in the junction between the NTD and the RBD domains. A mutation hot-spot in this junction is of interest as it can have a large effect to the transition between the closed and open states, the way how the RBD becomes exposed to ACE2. The site 316S is the only potential hot-spot mutation site in this junction that has been identified in the present study.

The RMSD values (17) are quite small so that the three clusters identified in the Figures 10 are a good match. The chain between sites 315-317 has a very similar topology to the chains 102-104 in Figures 7 and 286-288 in Figures 8. The clusters are also the same as in the case of 287D and therefore it is concluded that the site 316 is a good candidate for a S→A mutation hot-spot.

**Fig 10.**
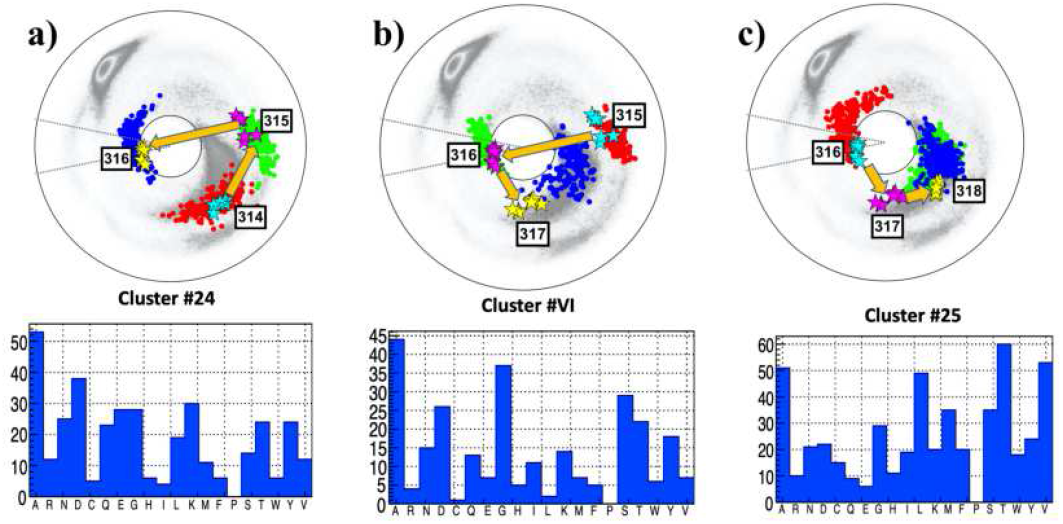
The (*κ_i_, τ_i_*) spectrum for *i* = 314 – 318; this engages sites 312 – 319 along the C*α* backbone. Panel a) The spectrum for P:(314, 315, 316), in the background of the best matching cluster #24 in [44]. The most common amino acids in the cluster is A, followed by D. Panel b) The spectrum for A:(315, 316, 317), in the background of the best matching cluster #VI in [44]. The most common amino acid in the cluster is A, followed by G. Panel c) The spectrum for F:(316, 317, 318), in the background of the best matching cluster #25 in [44]. The most common amino acid in the cluster are T and V, followed by A and L.

### The case of 464F

This site is located in the RBD domain of the subunit S1, and a mutation hot-spot can affect the binding to ACE2. Unfortunately, there are missing residues in the PDB data, only data for chain A in open state (PDB structure 6VYB) is available. The chain shown in Figures 11 is very similar to that in the case of 59F shown in Figures 6. The RMSD values are very low so that the three clusters are an excellent match. The statistical analysis shows that the site is a very good hot-spot candidate for a F→G mutation.

**Fig 11.**
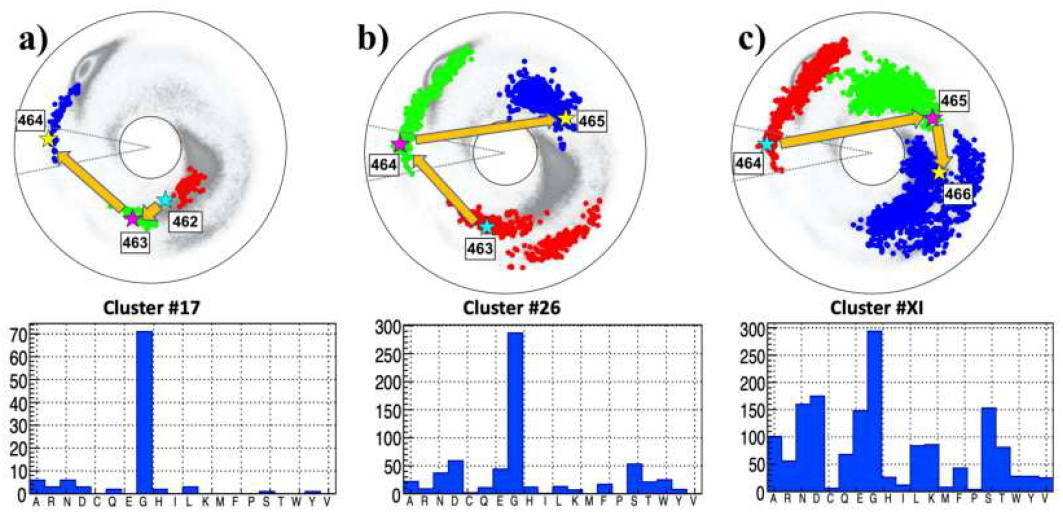
The (*κ_i_, τ_i_*) spectrum for *i* = 462 – 466; this engages sites 460 – 467 along the C*α* backbone. Panel a) The spectrum for P:(462, 463, 464), in the background of the best matching cluster #17 in [44]. Panel b) The spectrum for A:(463, 464, 465), in the background of the best matching cluster #26 in [44]. Panel c) The spectrum for F:(464, 465, 466), in the background of the best matching cluster #XI in [44]. The most common amino acid in all three clusters is G.

### The case of 1080A

According to the histogram in Figure 4 the fusion core between the HR1 and HR2 domains has a large number of sites that are proximal to a flattening point. However, the detailed investigation does not reveal any excellent candidate for a mutation hot-spot. An example is the site 1080A, the analysis results are summarized in Figures 12.

**Fig 12.**
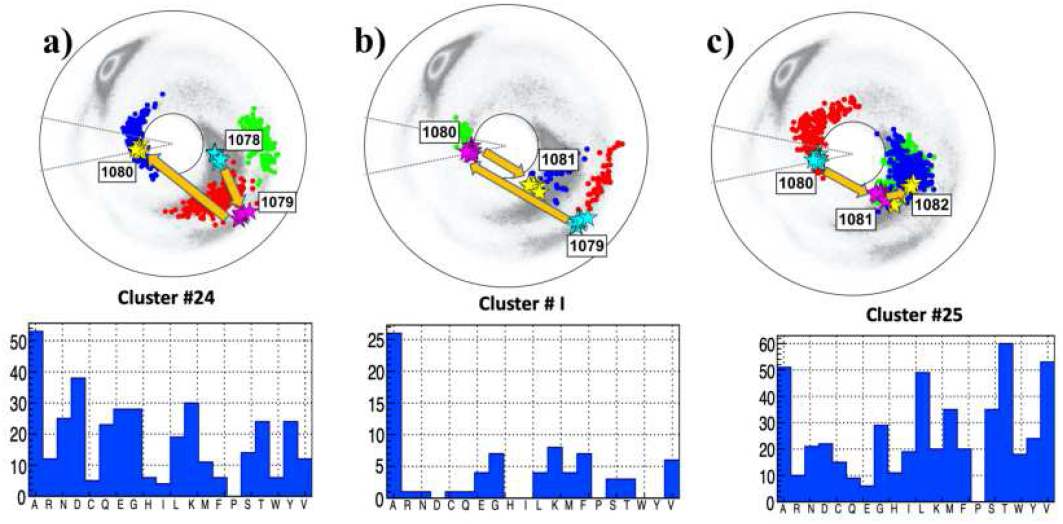
The (*κ_i_, τ_i_*) spectrum for *i* = 1078 – 1082; this engages sites 1076 – 1083 along the C*α* backbone. Panel a) The spectrum for P:(1078,1079.1080), in the background of the best matching cluster #24 in [44]. The most common amino acids in the cluster is A. Panel b) The spectrum for A:(1079,1080,1081), in the background of the best matching cluster #I in [44]. The most common amino acid in the cluster is A, followed by G. Panel c) The spectrum for F:(1080,1081,1082), in the background of the best matching cluster #25 in [44]. The most common amino acid in the cluster are T and V, followed by A and L.

The RMSD value for preceding cluster P:(1078,1079,1080) is not very good, but the value is excellent both for the adjacent A:(1079,1080,1081) and following F:(1080,1081,1082) clusters. Moreover, the chain shows that the site 1080 is an excellent candidate for a bi-flattening perestroika. But the statistical analysis reveals that the amino acid A, presently at this site, is also the most probable one and with very high probability in the case of the adjacent cluster. Thus, it appears that a recent mutation may have taken place at this site with the amino acid A as the substitution. The statistical analysis of the third cluster shows a relatively high propensity for T, V and L but all three are subject to steric hindrances and they also have a very low propensity in the adjacent cluster. The conclusion is that even though the vertex is proximal to a bi-flattening perestroika, it does not qualify as a mutation hot-spot. It is more likely that due to proximity of the bi-flattening perestroika, the site has exceptional flexibility with an important role in the fusion process. The residue 1080A can be a good target for the development of fusion inhibitors.

### The local topology of the N501Y mutation site

Thus far, in the present article, only those mutation hot-spot sites that are in the vicinity of a flattening point have been analyzed. However, the three local topologies in Figures 3 can give rise to various other kind of bifurcations as well. For this, the local topology of the recently observed new mutation at site 501N with N→Y substitution is now analyzed, for its bifurcation potential. Apparently, this is a substitution with considerable epidemiological consequences, it appears to cause increased transmissibility. For an epidemiological description see *e.g.* [31].

The Figures 13 describe the local topology of the site 501 prior to the N→Y substitution; data is presently available only for the chains B and C in the open state of 501N. In particular, no structural data is available for the mutated 501Y at this time. Except for the direction and the location of the end points, the topology of the trajectory in Figures 13 is akin that in Figure 3 c) with two crossings of the line of *β*-flattening point (but very near to the inflection point). There is no site in the vicinity of *τ* = 0 in Figures 13. Thus a bifurcation that relates to a flattening point, similar to that in the case of the D614G, appears unlikely.

**Fig 13.**
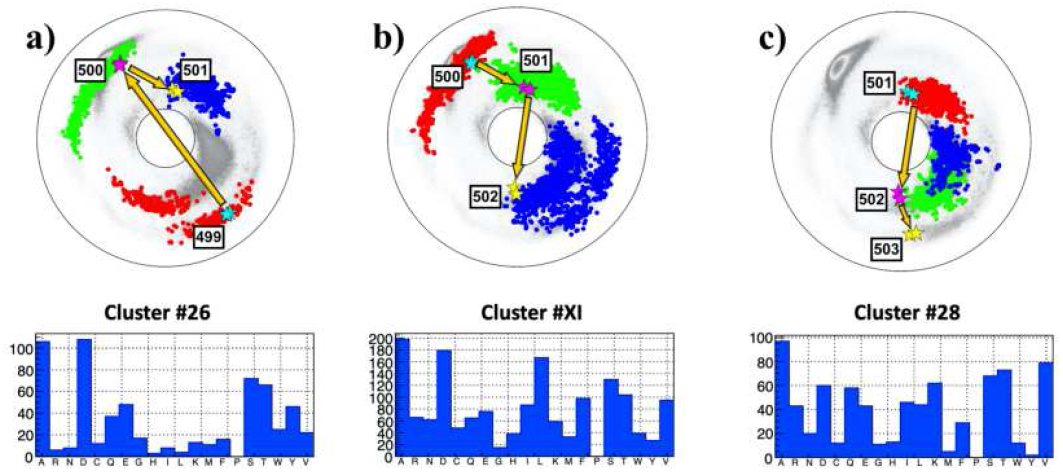
The (*κ_i_, τ_i_*) spectrum for *i* = 499 – 503; this engages sites 497 – 504 along the C*α* backbone. Data for 501N is only available in the open state chains B and C. Panel a) The spectrum for P:(499, 500, 501), in the background of the best matching cluster #26 in [44]. The most common amino acids in the cluster are A and D. Notably, Y is more common than N. Panel b) The spectrum for A:(500, 501, 502), in the background of the best matching cluster #XI in [44]. The most common amino acid in the cluster are A and D; now N is more common than Y. Panel c) The spectrum for F:(501,502, 503), in the background of the best matching cluster #28 in [44]. The most common amino acid in the cluster is A, there is also propensity to D but V is more common, N appears but Y is absent.

Notably both the segment connecting 499 to 500, and the segment connecting 501 to 502 pass very close to the inflection point *κ* = 0. This suggests that the local topology can be prone to a variant of an inflection point bifurcation that causes a transition either into the topology akin that in Figure 3 a) or that in Figure 3 b).

In Figures 13 a) and b) the RMSD values (17) are very small; in the case of Figure 13 a) the value is Δ_*min*_ =0.2 and in the case of Figure 13) the value is Δ_*min*_ = 0.16. Thus both spike protein segments are well represented by the ensuing clusters. But in the case of Figure 13 c) the value is much larger, Δ_*min*_ = 0.7 so that this segment is not well represented by the cluster. According to all three histograms in Figures 13 the likelihood of the observed N→Y substitution at site 501 should be lower than substitutions N→A or N→D by the similarly hydrophobic A and D. Furthermore, since Y has a much larger size than N, a priori the N→Y substitution requires an extensive conformational rearrangement which can be energetically costly. But a crude energetic Chimera [45] analysis of the neighborhood around the site 501N that is summarized in Figures 14 reveals that there is a “pocket” inside the spike protein structure that is large enough to accommodate Y with no major conformational re-arrangement; only a change in the orientation of 505Y is observed. Since the N501Y mutation is not in necessarily in contrast with stereochemical considerations, the energetic cost of a substitution can be minor.

**Fig 14.**
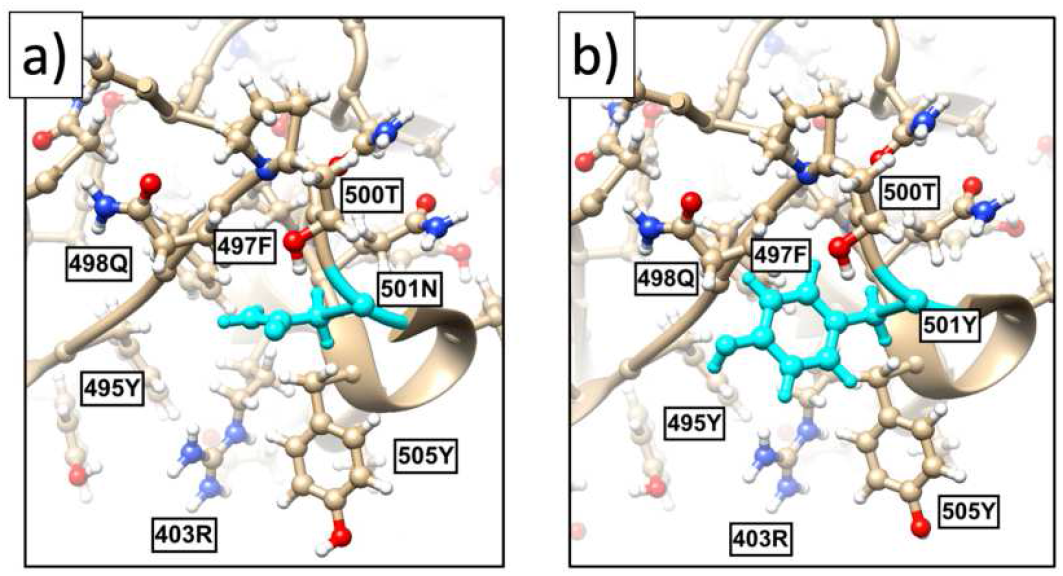
a) The present all-atom structure of the spike protein in the neighborhood of site 501. b) The effect of N→Y substitution as predicted by Chimera [45]. Besides a small change in the orientation of 505Y, despite the substantially larger size of Y in comparison to N there is no apparent steric hinder for the N→Y substitution at site 501.

The likely bifurcation that can accompany the N→Y substitution can be deduced by an analysis of all those fragments in the (Preceding) cluster #26 of Figure 13 a) where Y is in the third (Blue) position; this is the cluster where Y has the highest prevalence. The (Adjacent) cluster #XI can be analyzed similarly. But in the (Following) cluster #28 where the spike protein segment has a relatively low quality match, there is no Y.

This analysis starts with Figure 15 a): The initial (Red) dots of the fragments in the cluster #26 are naturally divided into two disjoint subsets. There are 42 backbone fragments that start in the subset labeled A in the Figure 15, and there are 4 fragments that start in the subset labeled B, with Y in the last (Blue) position in all fragments. The fragments in the subset A all pass very close the inflection point *κ* = 0, in a manner which is very similar to the 499-501 segment in Figure 13. The fragments in the subset B all cross the *τ* = 0 line in a manner that is topologically similar to the A-B-C segments in Figures 3 a) and b) (except for orientation). A comparison of the two subsets A and B in the Figure 15 b) reveals that there is a transition akin the peptide plane flip observed in [46]. For this, define the normal vector of the peptide plane between the C*α*(*i* – *i*) and C*α*(*i*), as the cross product between the vector **t**_*i*_ and the vector that points from C*α*(*i* – *i*) to the O(*i*) atom of the corresponding peptide plane. Similarly, define the normal vector of the peptide plane between C*α*(*i*) and C*α*(*i* + 1). Then, evaluate the angle between these two normal vectors. The result is shown in the histogram of Figure 15 b): For all entries in the subset A of Figure 15 a) the angle has a value which is very close to –*π*/2 while for all entries in the subset B the angle has a value that is very close to +*π*/2. The sign is determined by comparison to the virtual plane that is defined by C*α*(*i* – 1), C*α*(*i*) and C*α*(*i* + 1).

**Fig 15.**
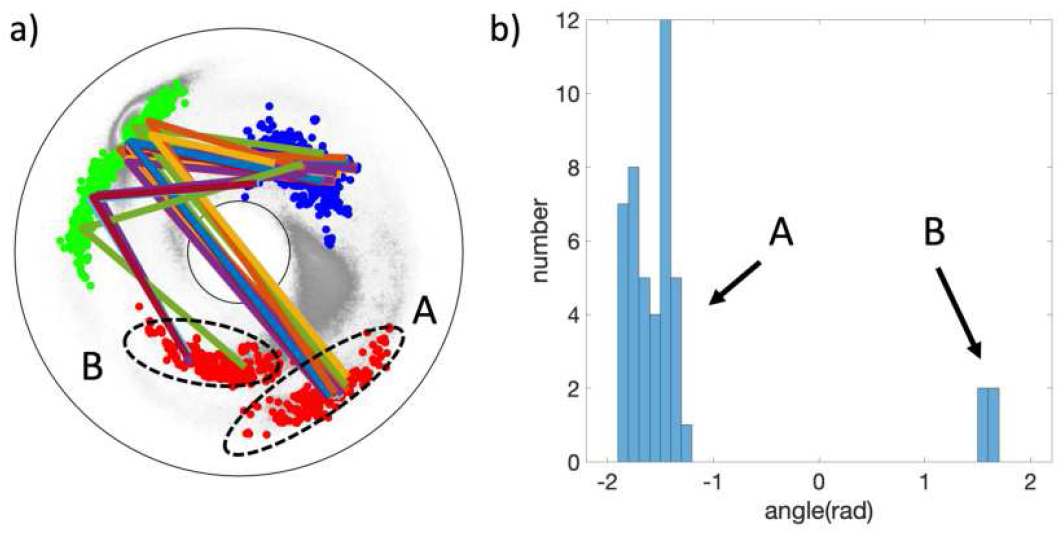
a) The fragments of cluster #26 where Y appears in the last (Blue) site can be divided into two disjoint subsets, labeled A and B. b) The subsets A and B differ by a ~ 180 degree rotation (flip) of the peptide plane between the second (Green) and third (Blue) site, around the connecting Frenet frame tangent vector **t**.

The present analysis proposes that the peptide plane between sites 500 and 501 of spike protein can be prone to an inflection point bifurcation where the peptide plane flips, corresponding to a rotation of ~ 180 degrees around the Frenet frame vector **t** between the ensuing C*α* atoms. Whether this bifurcation, or one that can occur between sites 501 and 502, plays a role in the transmissibility of the N501Y mutation can be determined once structural data becomes available.

Finally, there are recent mutations that have taken place in the spike protein with considerable epidemiological consequences, including the sites 484 and 677. Unfortunately, the structural data at and around these sites is presently missing in the available Protein Data Bank structures, thus the potential bifurcations that may be associated with these mutations can not yet be characterized.

## Conclusions

Topological techniques are commonly regarded to be among the most effective ones for addressing a wide range of physical problems. In particular in the case of a protein, where conformation is pivotal to biological function, topology should be a most valuable tool. However, thus far applications of topological methods in protein studies, and more generally in the study of filamental (bio)molecules such as DNA and RNA and other (bio)polymers, have been relatively sparse. The topological methodology that is developed here, for the purposes of protein studies, is designed to investigate bifurcations that can change the local topology of a space curve such as the protein backbone; similar studies on proteins have not been performed previously. The particular problem that is addressed here, is to try and pinpoint those hot-spot residues where a mutation can have the most profound conformational consequences: The hot-spot residues are those sites that are proximal to bifurcations where the local topology can change. The present methodology identifies these hot-spots using local topological considerations in combination with a comparative analysis that uses Protein Data Bank structures. But the methodology does not aim to actually predict, whether mutations at these hot-spot residues are more likely than at other sites. This needs to be deduced by other methods. The underlying mathematical structure is relatively new, it was developed mainly by Arnol’d starting late 1990’s. Prior to the present study it has thus far not found any (bio)physical applications. Thus the present study serves also as an invitation to apply these powerful methods of local topology to relate the structure and function of proteins, and to apply them to study the physical properties of biomolecules and filamental structures also more widely.

The methodology is developed using the spike protein of the SARS-CoV-2 virus as an example. Any other protein could be used equally but the choice of the spike protein is timely, reflecting the current pandemic situation. This choice also brings some downside, as the available structural data on the spike protein has been measured with relatively low resolution and is also partially lacking, at the time of writing: The Protein Data Bank structures that are used in the present study have been determined using electron microscopy with a resolution no better than around 2.8 – 3.5 Å. Missing residues in the available spike protein structures also limits the study. For example the recent epidemiologically important mutations that have been observed at the sites 484 and 677 could not be included in the study, due to lacking experimental data. The low resolution causes also some uncertainty in the proper identification of sites that are proximal to a flattening point. At the same time the statistical analysis that is used here for the identification of the pertinent PDB clusters is also preliminary, as it is based on a somewhat limited set of structural clusters that are measured with ultra high resolution. Therefore, a further development and refinement of the underlying statistical methodology is desired. Finally, the present article concerns only the local topology of a protein backbone. A more complete analysis should combine topological investigations with energetic studies. But that brings a practical limitation, as the presently available all-atom molecular force fields are still not very accurate and demand substantial computational resources. Here Chimera has been employed, as it is a method that can provide crude energetic estimates in the case complex proteins such as the spike protein, without undue need for computer resources.

## Acknowledgements

XP is supported by Beijing Institute of Technology Research Fund Program for Young Scholars. AJN is supported by the Carl Trygger Foundation Grant CTS 18:276, by the Swedish Research Council under Contract No. 2018-04411, and by COST Action CA17139. AJN is also partially supported by Grant No. 0657-2020-0015 of the Ministry of Science and Higher Education of Russia.

## References

1. Eguchi T, Gilkey PB, Hanson AJ. Gravitation, gauge theories and differential geometry. Physics reports. 1980; 66(6):213–393.

2. Eschrig H. Topology and geometry for physics. vol. 822. Springer Science & Business Media; 2011.

3. Nakahara M. Geometry, topology and physics. CRC Press; 2003.

4. Strogatz S. H. Nonlinear dynamics and chaos with with applications to physics, biology, chemistry, and engineering CRC Press; 2014

5. Flapan E, Wong H (Eds.) Topology and Geometry of Biopolymers AMS Contemporary Mathematics 2020;746

6. Arnold, V. I. (Ed.) (1994) Dynamical Systems V: Bifurcation Theory and Catastrophe Theory Encyclopaedia of Mathematical Sciences Vol. 5, Springer Verlag; 2014

7. Zhu N, Zhang D, Wang W, Li X, Yang B, Song J, et al. A novel coronavirus from patients with pneumonia in China, 2019. New England Journal of Medicine. 2020; 382:727–733

8. Lu R, Zhao X, Li J, Niu P, Yang B, Wu H, et al. Genomic characterisation and epidemiology of 2019 novel coronavirus: implications for virus origins and receptor binding. The Lancet. 2020;395(10224):565–574.

9. Andersen KG, Rambaut A, Lipkin WI, Holmes EC, Garry RF. The proximal origin of SARS-CoV-2. Nature medicine. 2020;26(4):450–452.

10. Benvenuto D, Giovanetti M, Ciccozzi A, Spoto S, Angeletti S, Ciccozzi M. The 2019-new coronavirus epidemic: evidence for virus evolution. Journal of medical virology. 2020;92(4):455–459.

11. Paraskevis D, Kostaki EG, Magiorkinis G, Panayiotakopoulos G, Sourvinos G, Tsiodras S. Full-genome evolutionary analysis of the novel corona virus (2019-nCoV) rejects the hypothesis of emergence as a result of a recent recombination event. Infection, Genetics and Evolution. 2020;79:104212.

12. Riou J, Althaus CL. Pattern of early human-to-human transmission of Wuhan 2019 novel coronavirus (2019-nCoV), December 2019 to January 2020. Eurosurveillance. 2020;25(4):2000058.

13. Zhao S, Lin Q, Ran J, Musa SS, Yang G, Wang W, et al. Preliminary estimation of the basic reproduction number of novel coronavirus (2019-nCoV) in China, from 2019 to 2020: A data-driven analysis in the early phase of the outbreak. International journal of infectious diseases. 2020;92:214–217.

14. Chan JFW, Yuan S, Kok KH, To KKW, Chu H, Yang J, et al. A familial cluster of pneumonia associated with the 2019 novel coronavirus indicating person-to-person transmission: a study of a family cluster. The Lancet. 2020;395(10223):514–523.

15. Phan LT, Nguyen TV, Luong QC, Nguyen TV, Nguyen HT, Le HQ, et al. Importation and human-to-human transmission of a novel coronavirus in Vietnam. New England Journal of Medicine. 2020;382(9):872–874.

16. Kissler SM, Tedijanto C, Goldstein E, Grad YH, Lipsitch M. Projecting the transmission dynamics of SARS-CoV-2 through the postpandemic period. Science. 2020;368(6493):860–868.

17. Wrapp D, Wang N, Corbett KS, Goldsmith JA, Hsieh CL, Abiona O, et al. Cryo-EM structure of the 2019-nCoV spike in the prefusion conformation. Science. 2020;367(6483):1260–1263.

18. Huang Y, Yang C, Xu Xf, Xu W, Liu Sw. Structural and functional properties of SARS-CoV-2 spike protein: potential antivirus drug development for COVID-19. Acta Pharmacologica Sinica. 2020;41:1141–1149

19. Walls AC, Park YJ, Tortorici MA, Wall A, McGuire AT, Veesler D. Structure, function, and antigenicity of the SARS-CoV-2 spike glycoprotein. Cell. 2020;181(2):281–292

20. Dai W, Zhang B, Jiang XM, Su H, Li J, Zhao Y, et al. Structure-based design of antiviral drug candidates targeting the SARS-CoV-2 main protease. Science. 2020;368(6497):1331–1335.

21. Gordon DE, Hiatt J, Bouhaddou M, Rezelj VV, Ulferts S, Braberg H, et al. Comparative host-coronavirus protein interaction networks reveal pan-viral disease mechanisms. Science. 2020;370(6521):eabe9403

22. Li JY, Liao CH, Wang Q, Tan YJ, Luo R, Qiu Y, et al. The ORF6, ORF8 and nucleocapsid proteins of SARS-CoV-2 inhibit type I interferon signaling pathway. Virus research. 2020;286:198074.

23. Gordon DE, Jang GM, Bouhaddou M, Xu J, Obernier K, White KM, et al. A SARS-CoV-2 protein interaction map reveals targets for drug repurposing. Nature. 2020; 583:459–468

24. Hyeonuk W, Sang-Jun P, Yeol KC, Taeyong P, Maham T, Yiwei C, et al. Developing a Fully-glycosylated Full-length SARS-CoV-2 Spike Protein Model in a Viral Membrane. The journal of physical chemistry B. 2020;124:7128–7137.

25. Chi X, Yan R, Zhang J, Zhang G, Zhang Y, Hao M, et al. A neutralizing human antibody binds to the N-terminal domain of the Spike protein of SARS-CoV-2. Science. 2020;369(6504):650–655.

26. Letko M, Marzi A, Munster V. Functional assessment of cell entry and receptor usage for SARS-CoV-2 and other lineage B betacoronaviruses. Nature microbiology. 2020;5(4):562–569.

27. Hoffmann M, Kleine-Weber H, Pöhlmann S. A multibasic cleavage site in the spike protein of SARS-CoV-2 is essential for infection of human lung cells. Molecular Cell. 2020;78(4):779–784.e5

28. Shang J, Wan Y, Luo C, Ye G, Geng Q, Auerbach A, et al. Cell entry mechanisms of SARS-CoV-2. Proceedings of the National Academy of Sciences. 2020;117(21):11727–11734.

29. Hoffmann M, Kleine-Weber H, Schroeder S, Krüger N, Herrler T, Erichsen S, et al. SARS-CoV-2 cell entry depends on ACE2 and TMPRSS2 and is blocked by a clinically proven protease inhibitor. Cell. 2020;181(2):271–280.e8

30. Korber B, Fischer WM, Gnanakaran S, Yoon H, Theiler J, Abfalterer W, et al. Tracking changes in SARS-CoV-2 Spike: evidence that D614G increases infectivity of the COVID-19 virus. Cell. 2020;182(4):812–827.

31. Mahase E. Covid-19: What have we learnt about the new variant in the UK? BMJ. 2020;371. doi:10.1136/bmj.m4944.

32. Hodcroft E. B. et.al Emergence in late 2020 of multiple lineages of SARS-CoV-2 Spike protein variants affecting amino acid position 677. medRxiv 2021.02.12.21251658

33. Arnold V. Singularities of caustics and wave fronts. vol. 62. Springer Science & Business Media; 2013.

34. Arnold V. The geometry of spherical curves and the algebra of quaternions. Russian Mathematical Surveys. 1995;50(1):1.

35. Arnold V. On the Number of Flattening Point on Space Curves. AMS Trans, Ser. 1996;171:11.

36. Aicardi F. Self-linking of spatial curves without inflections and its applications. Functional Analysis and Its Applications. 2000;34(2):79–85.

37. Uribe-Vargas R. On singularities,” perestroikas” and differential geometry of space curves. Enseignement Mathematique. 2004;50(1/2):69–102.

38. Yurkovetskiy L, Wang X, Pascal KE, Tomkins-Tinch C, Nyalile TP, Wang Y, et al. Structural and functional analysis of the D614G SARS-CoV-2 spike protein variant. Cell. 2020;183(3):739–751.e8

39. Spivak MD. A comprehensive introduction to differential geometry. vol. 5. Publish or perish; 1970.

40. Hu S, Lundgren M, Niemi AJ. Discrete Frenet frame, inflection point solitons, and curve visualization with applications to folded proteins. Physical Review E. 2011;83(6):061908.

41. Lundgren M, Niemi AJ, Sha F. Protein loops, solitons, and side-chain visualization with applications to the left-handed helix region. Physical Review E. 2012;85(6):061909.

42. Lundgren M, Krokhotin A, Niemi AJ. Topology and structural self-organization in folded proteins. Physical Review E. 2013;88(4):042709.

43. Hinsen K, Hu S, Kneller GR, Niemi AJ. A comparison of reduced coordinate sets for describing protein structure. The Journal of Chemical Physics. 2013;139(12):124115.

44. Peng X, He J, Niemi AJ. Clustering and percolation in protein loop structures. BMC structural biology. 2015;15(1):22.

45. Shapovalov MV, Dunbrack Jr RL. A smoothed backbone-dependent rotamer library for proteins derived from adaptive kernel density estimates and regressions. Structure. 2011;19(6):844–858.

46. Liu J, Dai J, He J, Peng X, Niemi AJ. Can the geometry of all-atom protein trajectories be reconstructed from the knowledge of C α time evolution? A study of peptide plane O and side chain C *β* atoms. The Journal of chemical physics. 2019;150(22):225103.

